# Hydroxychloroquine increased psychiatric-like behaviors and disrupted the expression of related genes in the mouse brain

**DOI:** 10.1101/2020.09.27.316158

**Authors:** H Xu, XY Zhang, WW Wang, JS Wang

**Author notes:** **Corresponding Author**: Wang Jiesi, PhD, Institute of Psychology, Chinese Academy of Sciences, 16 Lincui Road, Chaoyang District, Beijing 100101, China.

## Abstract

Hydroxychloroquine (HCQ), which has been proposed as a therapeutic or prophylactic drug for SARS-COV-2, has been administered to thousands of individuals with varying efficacy; however, our understanding of its adverse effects is insufficient. It was reported that HCQ induced psychiatric symptoms in a few patients with autoimmune diseases, but it is still uncertain whether HCQ poses a risk to mental health. Therefore, in this study, we treated healthy mice with two different doses of HCQ that are comparable to clinically administered doses for 7 days. Psychiatric-like behaviors and the expression of related molecules in the brain were evaluated at two time points, i.e., 24 h and 10 days after drug administration. We found that HCQ increased anxiety behavior at both 24 h and 10 days and enhanced depressive behavior at 24 h. Furthermore, HCQ decreased the mRNA expression of interleukin-1beta and corticotropin-releasing hormone (Crh) in the hippocampus and decreased the mRNA expression of brain-derived neurotrophic factor (Bdnf) in both the hippocampus and amygdala. Most of these behavioral and molecular changes were sustained beyond 10 days after drug administration, and some of them were dose-dependent. Although this animal study does not prove that HCQ has a similar effect in humans, it indicates that HCQ poses a significant risk to mental health and suggests that further clinical investigation is essential. According to our data, we recommend that HCQ be carefully used as a prophylactic drug in people who are susceptible to mental disorders.

## Introduction

At present, tens of millions of people worldwide have been infected with coronavirus disease 2019 (SARS-COV-2). Among the many potential therapeutic drugs for SARS-COV-2, society has perhaps paid the most attention to hydroxychloroquine (HCQ). Although emerging evidence does not suggest that it is efficacious in treating SARS-COV-2 (1–3), it has already been administered to thousands of individuals. Furthermore, more than hundred clinical trials of HCQ involving tens of thousands of potential participants have been registered. HCQ is not only being used to treat patients with SARS-COV-2 but is also being used as a pre-exposure or postexposure prophylactic drug for SARS-COV-2 in healthy populations who are at high risk of exposure to SARS-COV-2 (4, 5). Although HCQ has been used clinically for decades to treat malaria and autoimmune diseases, such as systemic lupus erythematosus (SLE), Sjögren’s syndrome (SS) and rheumatoid arthritis (RA) (6), our understanding of its potential adverse effects, especially in patients with SARS-COV-2 and uninfected individuals, is insufficient.

There have been reports of serious adverse effects related to HCQ, including retinopathy, QT interval prolongation, hypoglycemia, cardiomyopathy, neuropsychiatric effects and so on (7, 8), and QT interval prolongation is the most closely monitored adverse effect in patients with SARS-COV-2. Less attention has been paid to the effects of HCQ on mental health, although multiple studies have reported severe psychiatric symptoms, including agitation, hallucinations, anxiety, depression, and suicidal ideation, in patients treated with HCQ or chloroquine (9–12). However, whether these neuropsychiatric symptoms are caused by HCQ alone or by the interaction between HCQ and the disease is not well studied, possibly due to the limited number of reported cases. Therefore, whether HCQ poses a serious risk to mental health is still not clear.

In addition to acting as an antimalaria drug, HCQ is an immunomodulator that exerts broad anti-inflammatory effects, including inhibition of antigen presentation to dendritic cells, reduced signaling of both B and T cells and reduced cytokine production in macrophages (6, 13). Microglia, as macrophages that reside in the brain and play critical roles in neuronal functions, are also potential targets of HCQ (14). However, the ability of HCQ to cross the blood-brain barrier (BBB) is quite limited (15, 16); however, considering that HCQ has a very long half-life in vivo (6), its direct effect on microglia should not be underestimated. In addition, HCQ may affect the CNS via its inhibitory on peripheral cytokines, such as IL-1β or TNF-α, which can bind to the vascular endothelium of the brain or directly entering the CNS (17), resulting in a reduction in the production of corresponding cerebral cytokines and induction of a neuroinflammatory deficit in the brain. Although hypo-neuroinflammation is not as concerning as hyper-neuroinflammation, and the latter has been indicated as a potential pathology of multiple psychiatric disorders (18), emerging evidence suggests that maintaining the balance of the immune system is more important than preventing inflammation in the CNS (19) and that low levels of inflammation in the brain can also have negative effects on cognition and behaviors (20, 21). Therefore, the immunosuppressive effect of HCQ may have negative effects on the CNS, especially in healthy individuals who do not suffer from hyperinflammation induced by infection.

Therefore, in this study, the potential effects of HCQ on mental health were investigated in healthy mice. The animals were administered HCQ for 7 days, and psychiatric-like behaviors related to anxiety, depression, and cognitive impairment were evaluated. Furthermore, the expression of genes, including brain-derived neurotrophic factor (Bdnf), corticotropin-releasing hormone (Crh) and interleukin-1beta (IL-1β), which are closely related to mental health, were also measured in two brain areas, namely, the hippocampus and amygdala, which play critical roles in mental health and in which cytokines are produced notably(17). In this study, we calculated appropriate doses of HCQ for mice based on clinically administered doses of HCQ according to the average body surface area of humans and mice. The clinical dosages used for the calculations were 800 and 400 mg on the first day followed by 400 and 200 mg daily for the next 6 days. We evaluated behavior and gene expression at two time points, i.e., 24 h and 10 days after drug administration, to evaluate the immediate and lasting effects of the drug.

## Methods

### Animals

The male C57BL/6J mice used in this study were purchased from Vital River Laboratory Animal Center (Beijing, China) at the age of 8 weeks. After a week of adaptation, the mice were randomly divided into two groups: the immediate group and the lasting group. The mice in each group were further randomly divided into three subgroups, i.e., the high-dose, low-dose and vehicle groups; there were 7~8 animals per subgroup. All animals were maintained under controlled environmental conditions (12 h light/12 h dark cycle, lights on at 7 am; ambient temperature 22-24 °C; humidity 55 ± 10%) with free access to food and water. All procedures were approved by the Institutional Review Board of the Institute of Psychology, Chinese Academy of Sciences and performed in compliance with the National Institutes of Health Guide for the Care and Use of Laboratory Animals.

### Drug Administration

Hydroxychloroquine sulfate (Ark Pharm, Chicago, USA) was dissolved in sterile phosphate-buffered saline (PBS), pH=7.4. The drug was intraperitoneally (i.p.) injected twice per day (every 12 hours) for 7 consecutive days. On the first day, the high dose of HCQ was 82 mg/kg/injection, and the low dose was 41 mg/kg/injection; HCG was given at half of these doses for the next 6 days. The body weight of each mouse was recorded every two days during administration and every five days after administration.

### Behavioral Tests

To test the immediate/lasting effects of HCQ on behavior and the expression of molecules in the brain, behavioral tests were performed 24 h and 10 days after the last injection. The interval between each test was 24 h. For each test, the apparatus was cleaned with 75% ethanol to eliminate the influence of excrement or odor on subsequent animals. The tests were performed in the following order.

### Novel Object Recognition (NOR) Test

The NOR test was used to evaluate the locomotor activity and working memory of the mice. The test consisted of three phases, including the adaptation phase (Phase 1, 30 min), the training phase (Phase 2, 10 min) and the test phase (Phase 3, 10 min). In Phase 1, the mice were placed in the apparatus (L*W*H: 30 cm*30 cm*40 cm, with dim light), allowed to explore freely for 30 min and then returned to their home cages. During the exploration period in Phase 1, the trajectory of each mouse was captured with a camera and automatically tracked and recorded with software (Anilab, Suzhou, China). The distance traveled every 5 min throughout the 30-min period was recorded as a measure of the locomotor activity of the mice. In Phase 2, two identical objects (Lego bricks, 3 cm*3 cm*6 cm) were placed in opposite corners of the apparatus 6 cm from either side of the wall. The mice were introduced from the corner not containing an object, allowed to explore for 10 min and then returned to their home cages for 1 h. In Phase 3, one of the two objects were replaced with a novel object of similar size but of a different color and shape. The mice were then placed in the apparatus again and allowed to explore for an additional 10 min. The exploration of each mice was recorded with an overhead camera and then hand-scored by a pretrained independent investigator blinded to the experimental groups. The total sniff time and the discrimination index ((sniff time for novel object-sniff time for familiar object)/(total sniff time)) (22) were the main parameters measured in this test. Discrimination index data for a few animals that showed a lack of exploratory activity (<1 s for one object) were excluded.

### Elevated plus-maze (EPM) test

The day after the locomotor and novel object recognition tests were performed, the mice were subjected to the EPM test in a quiet dim room to evaluate anxiety-like behaviors. The mice were placed in the center of an elevated maze facing the open arms at the beginning of the test. During the test, the mice were allowed to freely explore the maze for 10 min, and the movement trajectory and time spent in each area were automatically recorded by software. The duration of time spent in the open arms and in the closed arms was reported.

### Forced swim test (FST)

The FST was performed 24 h after the EPM test to measure depression-like behavior. The mice were forced to swim individually for 6 min in a glass cylinder with a diameter of 15 cm filled 20 cm deep with water at a temperature of 25±1 °C. Swimming behavior was recorded using a horizontal video camera, and immobility time (during which the mice exhibited no movement except for that of the whiskers) was hand-scored by two independent observers blinded to the experimental groups. The immobility time of each mouse was calculated as the average of the times measured by the two observers.

### RNA extraction and Q-PCR

The mice were sacrificed 24 h after all the behavioral tests were completed. The hippocampus and amygdala were quickly extracted from each brain by using a mouse brain slicer. The tissues were immediately frozen on dry ice and then stored at −80 °C. RNA was extracted from the tissues by using the RNAprep Pure Tissue Kit (Tiangen, Beijing, China) according to the standard protocol. The RNA concentration was quantified using the NanoDrop system (Thermo Fisher Scientific, MA, USA), and the quality of the RNA was evaluated by gel electrophoresis. One microgram of total RNA from each sample was reverse transcribed into first-strand cDNA by using the M-MLV kit (Promega Corporation, Madison, WI, USA) following the manufacturer’s protocol. mRNA expression levels were determined by real-time PCR using SYBR™ Green PCR Master Mix (Life Technologies, California, USA) on an ABI 7500 system, and dissociation curve analysis was performed at the end of each run. Each sample was individually amplified and duplicated, and then the average value was used for subsequent analyses. Relative mRNA expression was calculated by the standard 2–ΔΔCt method using glyceraldehyde 3-phosphate dehydrogenase (Gapdh) as a housekeeping gene.

The following forward (F) and reverse (R) primers (BGI Tech, Beijing, China) were used:

Il-1β: forward: TGTCTGAAGCAGCTATGGCAAC; reverse: CTGCCTGAAGCTCTTGTTGATG;
Bdnf: forward: TGGCTGACACTTTTGAGCAC; reverse: AAGTGTACAAGTCCGCGTCC;
Crh: forward: GATCTCACCTTCCACCTTCTG; reverse: CGCAACATTTCATTTCCCGA;
Gapdh: forward: CAAGCTCATTTCCTGGTATGAC; reverse: CTGGGATGGAAATTGTGAGG.

### Statistical Analysis

The data are presented as the mean ± SEM and were analyzed by GraphPad 8.0. Comparisons between the three groups were performed using one-way or two-way ANOVA, and LSD *post hoc* multiple comparisons were carried out. Statistical significance for all analyses was set at *p*⩽0.05.

## Results

### Effects of HCQ on body weight and locomotor activity

As shown in figure 1A, two-way ANOVA indicated that there was a significant effect of time (F(2.461, 49.23) = 101.1, P<0.0001) and a significant time × drug interaction effect (F(10, 100) = 2.668, P=0.0062) but no significant effect of drug treatment (F(2, 20) = 2.723, P=0.0899). Post hoc tests indicated that the rate of increase in body weight, which was significantly lower in the low-dose group (P=0.0479) and high-dose group (P=0.0155) than in the PBS group, was significantly different between groups 24 h after injection, but the difference between the low-dose group and high-dose group was not significant (P=0.1925). The rate of increase in weight over subsequent days were not significant.

**Figure 1:**
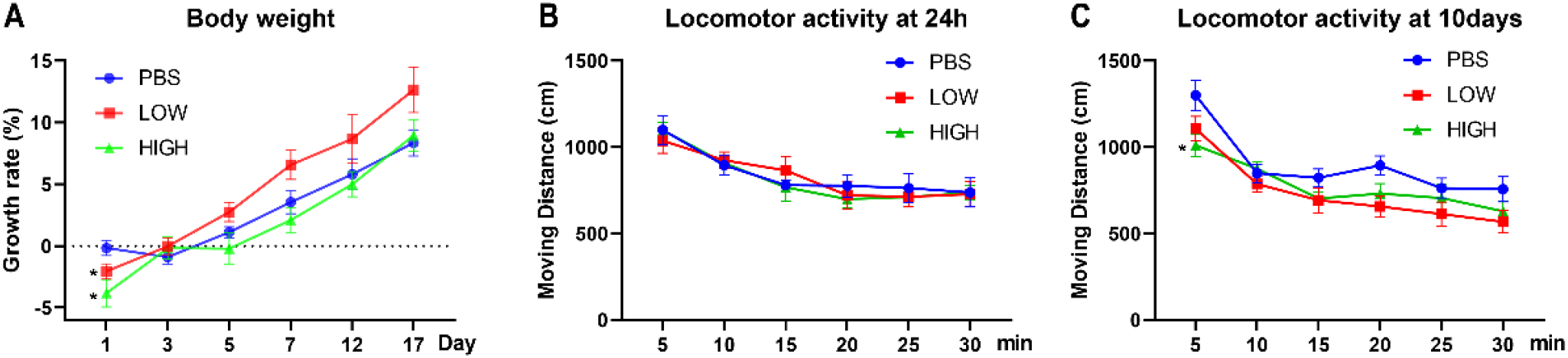
Effects of HCQ on body weight and locomotor activity. (A) The rate of increase in body weight of mice in the lasting group. (B) The locomotor activity of mice in the immediate group. (C) The locomotor activity of mice in the lasting group (n=7-8 for each subgroup; two-way ANOVA, \**p*<0.05 compared with the PBS group).

For locomotor activity (figure 1B, C), two-way ANOVA indicated that there was a significant effect of time (immediate: F(3.418, 75.20) = 25.86, P<0.0001; lasting: F(2.370, 47.40) = 39.25, P<0.0001) but no interaction effect (immediate: F(10, 110) = 0.5253, P=0.8691; lasting: F(10, 100) = 1.405, P=0.1891) in both the immediate and lasting groups. The drug effect was borderline significant in the lasting test (F(2, 20) = 3.488, P=0.0502) but not in the immediate test (F(2, 22) = 0.05869, P=0.9431). Post hoc tests indicated that the locomotor activity in the first 5 min was significantly lower in the high-dose group than in the PBS group (P =0.0215) in the lasting group.

### Immediate effects of HCQ on behavior

Figure 2 reveals the effect of HCQ on anxiety-like behavior, novel object recognition ability, and depressive-like behavior.

**Figure 2:**
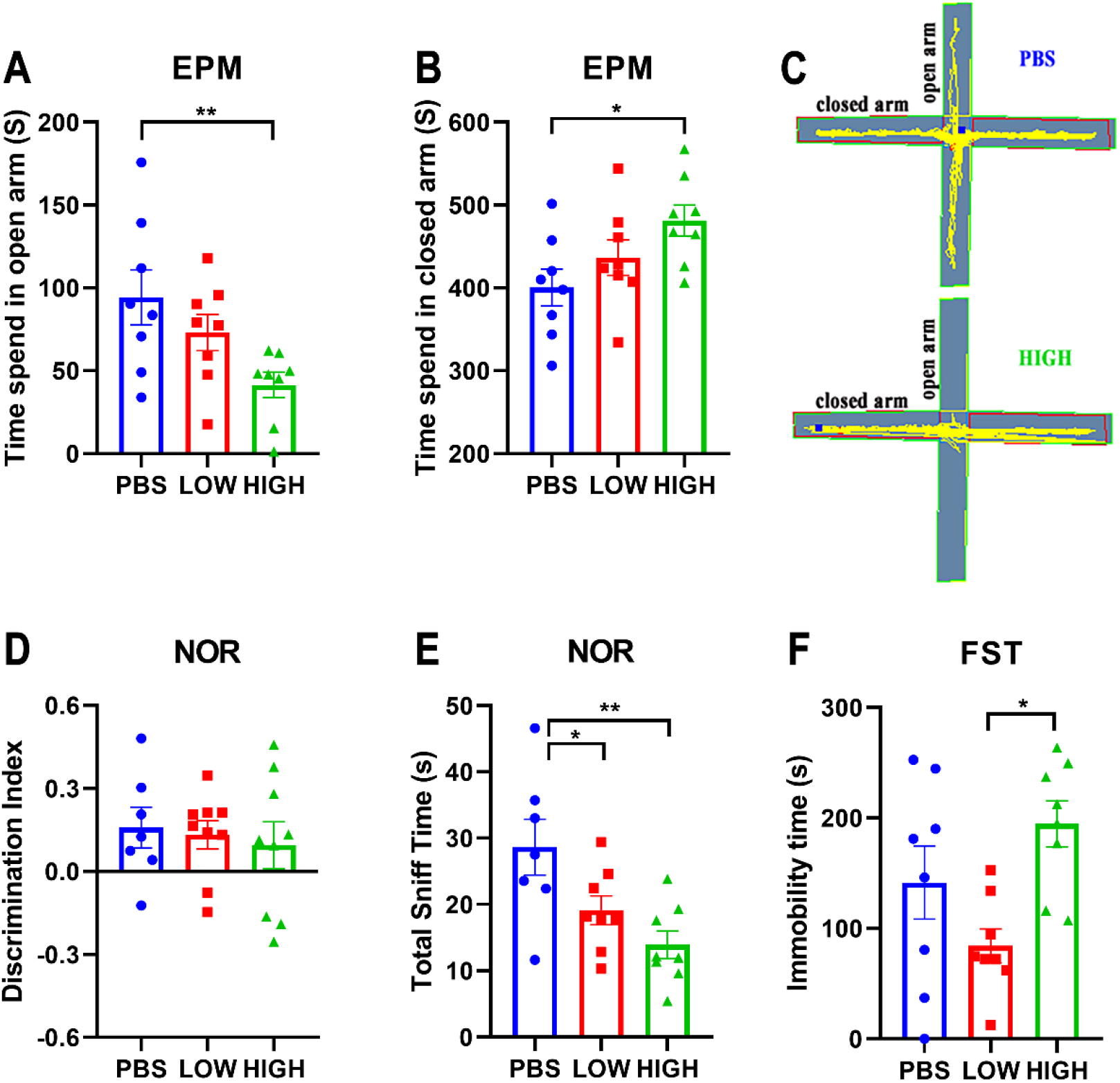
Effects of HCQ on the behaviors of mice in the immediate group. (A-C) Elevated plus-maze test. (A) Time spent in the open arms. (B) Time spent in the closed arms. (C) Typical trajectory of mice in the high-dose group and PBS group. (D&E) The novel object recognition test. (D) The discrimination index ((novel-familiar)/(novel+familiar)). (E) The total time spent sniffing. (F) The immobility time in the forced swim test (n=8 for each subgroup; one-way ANOVA; \**p*<0.05, ** *p*<0.01).

In the EPM test (figure 2A, B). One-way ANOVA indicated that there was a significant effect of HCQ treatment on the time spent in the open arms (F(2, 21) = 4.706, p=0.0205) and closed arms (F(2, 21) = 3.773, p=0.0398) at 48 h after the final injection. Post hoc analysis showed that the mice in the high-dose group spent much more time in the closed arms (481.3±18.73 vs. 400.6 ±22.00, p=0.0122) and less time in the open arms (41.35±7.65 vs. 94.33 vs. 16.55, p=0.0061) than mice in the PBS group.

For the novel object recognition test (figure 2D, E), one-way ANOVA indicated that there was no significant effect of HCQ treatment on the discrimination index at 24 h after the final injection (F(2, 22) = 0.1973, P=0.8224). However, there was a significant difference in total sniff time (F(2, 20) = 6.528, P=0.0066). Post hoc analysis revealed that mice in the high-dose group and low-dose group spent less time sniffing (13.93±2.092 vs. 28.64±4.219, p=0.001, and 19.13±2.188 vs. 28.64±4.219, p=0.0314, respectively) than mice in the PBS group.

For the FST (figure 2F), one-way ANOVA indicated that there was a significant effect of HCQ treatment on the immobility time at 96 h after the final injection (F(2, 21) = 5.185, P=0.0148). Post hoc analysis showed that the mice in the high-dose group exhibited a much longer immobility time than the mice in the low-dose group (194.7±20.72 vs. 84.25±15.4, p=0.0110).

### Lasting effects of HCQ on behavior

Figure 3 reveals the lasting effect of HCQ on anxiety-like behavior, depressive-like behavior, and novel object recognition ability.

**Figure 3:**
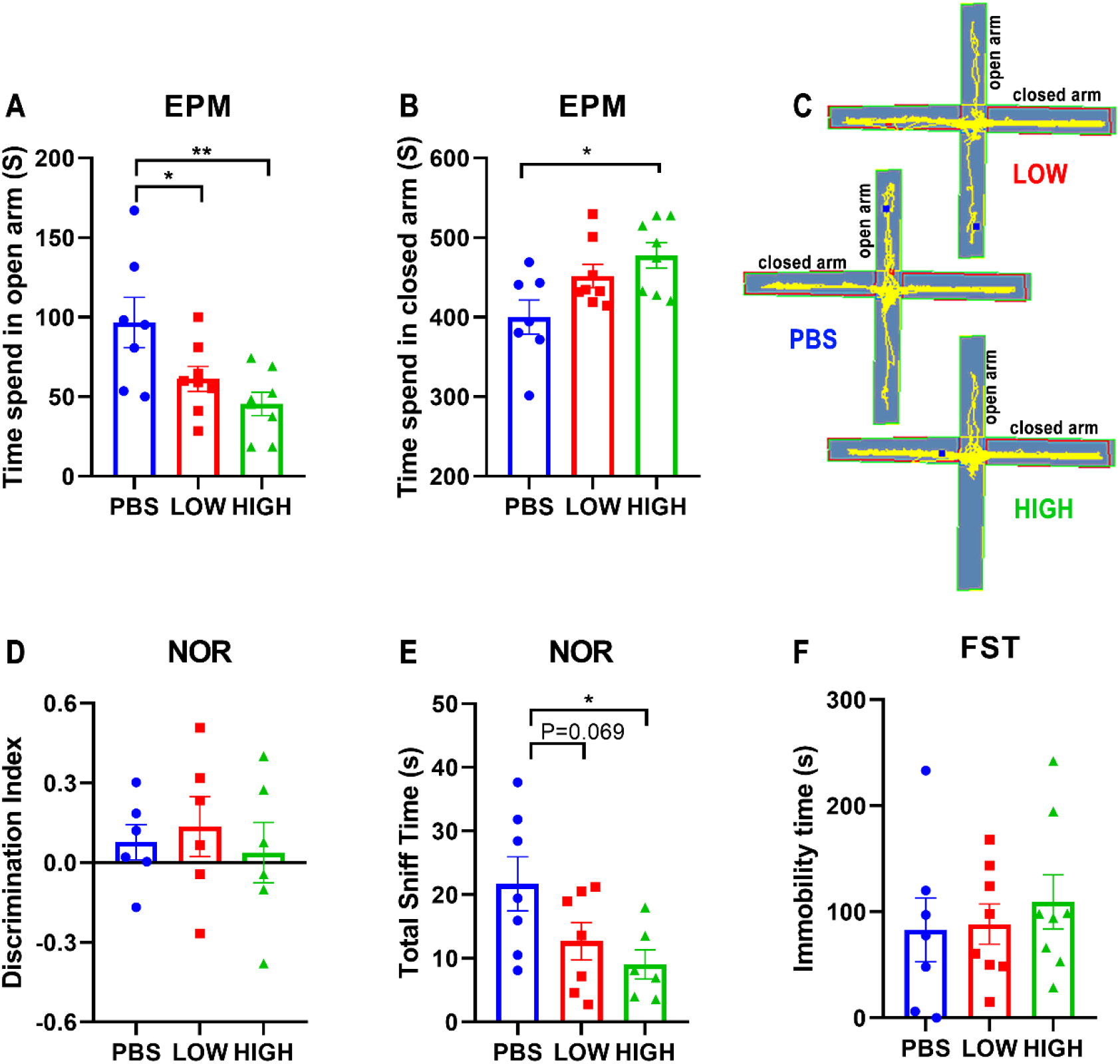
Effects of HCQ on the behavior of mice in the lasting group. (A-C) Elevated plus-maze test. (A) Time spent in the open arms. (B) Time spent in the closed arms. (C) Typical trajectory of mice in the high-dose, low-dose and PBS groups. (D&E) The novel object recognition test. (D) The discrimination index ((novel-familiar)/(novel+familiar)). (E) The total time spent sniffing. (F) The immobility time in the forced swim test (n=7-8 for each subgroup; one-way ANOVA; \**p*<0.05, ** *p*<0.01).

For the EPM test (figure 4A, B), one-way ANOVA indicated that there was a significant lasting effect of HCQ treatment on the time spent in the open arms (F(2, 20) = 5.982, p=0.0092) and closed arms (F(2, 20) = 5.026, p=0.017) at 12 days after the final injection. Post hoc analysis showed that the mice in the high-dose group spent much more time in the closed arms (477.7±16.06 vs. 400.2±21.39, p=0.0139) than the mice in the PBS group. Mice in the low-dose group and high-dose group spent less time in the open arms than mice in the PBS group (61.18±7.86 vs. 96.68± 15.79, p<0.05, and 45.55±7.29 vs. 96.68±15.79, p=0.0029, respectively) Anxiety-like behavior seemed to become more obvious over time.

**Figure 4:**
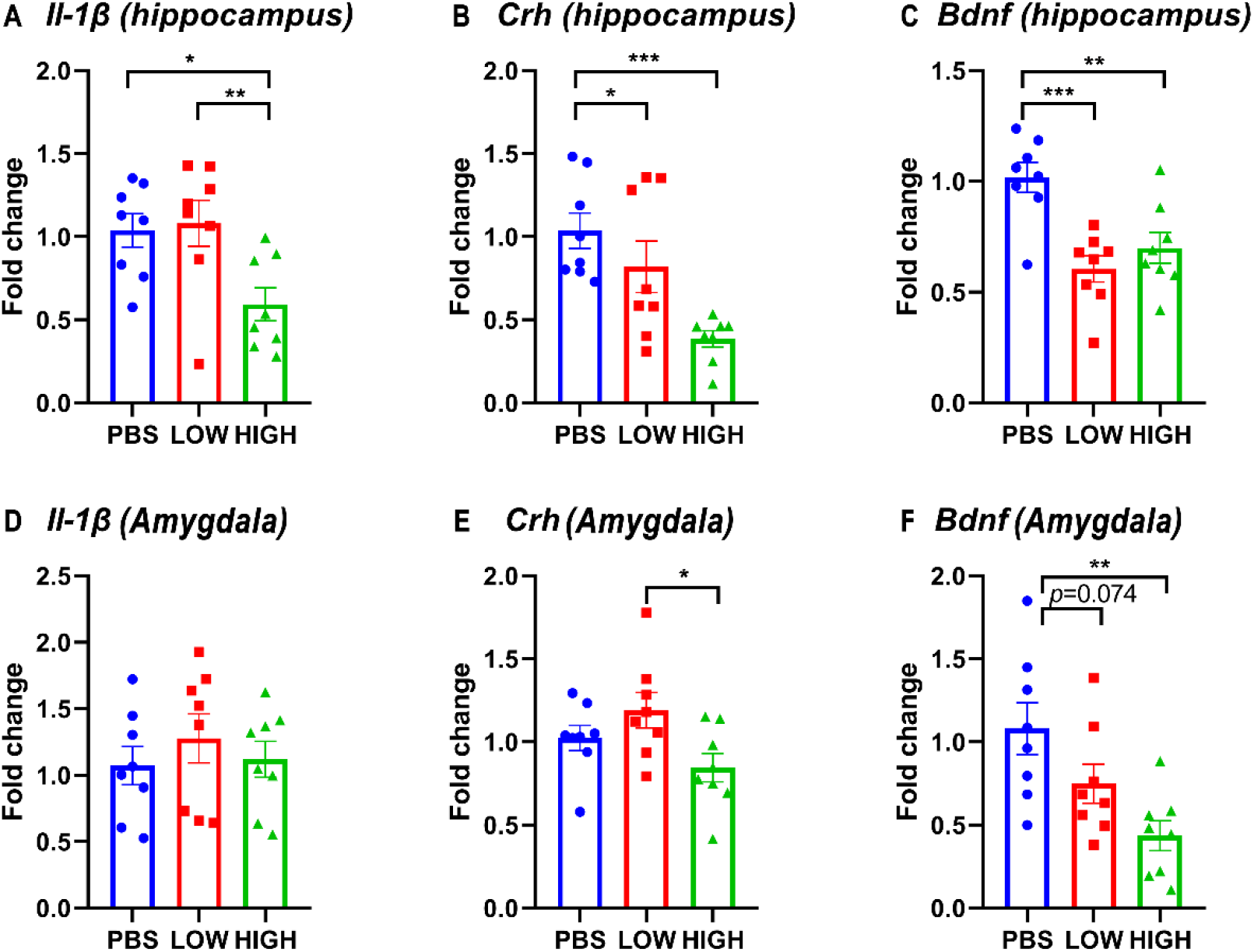
Effects of HCQ on the mRNA expression of genes in the hippocampus and amygdala of mice in the immediate group. (A-C) mRNA expression of (A) Il-1β, (B) Crh, and (C) Bdnf in the hippocampus. (D-F) mRNA expression of (D) Il-1β, (E) Crh, and (F) Bdnf in the amygdala (n=8 for each subgroup; one-way ANOVA; \**p*<0.05, ** *p*<0.01, *** *p*<0.005).

**Figure 5:**
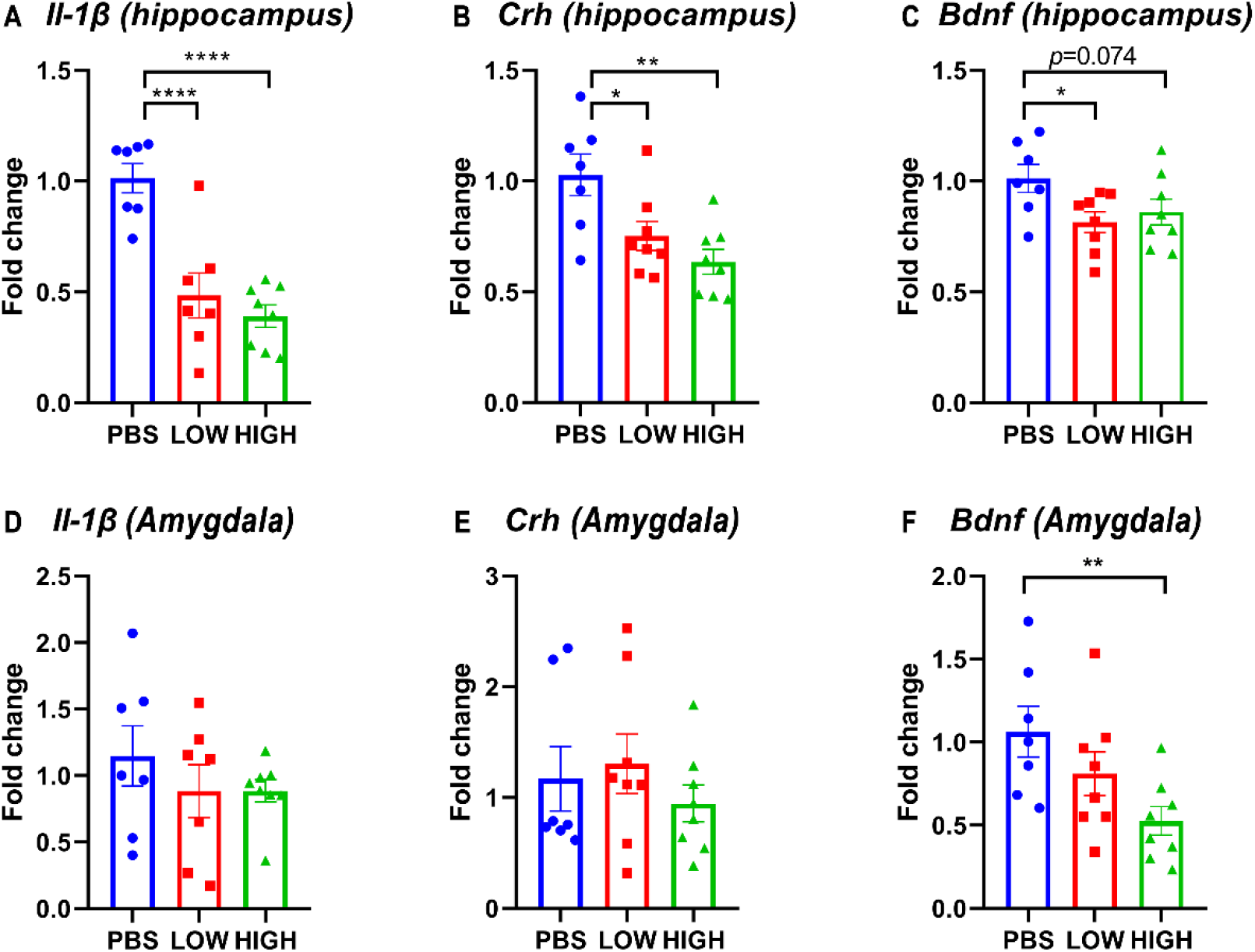
Effects of HCQ on the mRNA expression of genes in the hippocampus and amygdala of mice in the lasting group. (A-C) mRNA expression of (A) Il-1β, (B) Crh, and (C) Bdnf in the hippocampus. (D-F) mRNA expression of (D) Il-1β, (E) Crh, and (F) Bdnf in the amygdala (n=7-8 for each subgroup; one-way ANOVA; \**p*<0.05, ** *p*<0.01, \*\*\**p*<0.005).

For the novel object recognition test (figure 4D, E), one-way ANOVA indicated that there was no significant effect of HCQ treatment on the discrimination index at 10 days after the final injection (F(2, 15) = 0.2426, P= 0.7876). However, there was a significant difference in total sniff time (F(2, 17) = 3.754, P=0.0446). Post hoc analysis revealed that mice in the high-dose group spent less time sniffing (9.023±2.314 vs. 21.70±4.226, p=0.0176) than mice in the PBS group. For the FST (figure 4F), one-way ANOVA indicated that there was no significant effect of HCQ treatment on immobility time at 13 days after the final injection (F(2, 19) = 1.243, P=0.3109).

### Immediate effects of HCQ on the mRNA expression of Bdnf, Il-1β, and Crh in the hippocampus and amygdala

As shown in figure 4, HCQ rapidly altered the mRNA expression levels of Bdnf, Il-1β, and Crh in the hippocampus and amygdala. One-way ANOVA indicated that there was a significant effect of drug treatment on the expression of Il-1 β in the hippocampus (F(2, 21) = 5.635, P=0.0110) but not in the amygdala (F(2, 21) = 0.4723, P=0.6300). Post hoc analysis revealed that the mRNA expression of IL1-β in the high-dose group was significantly lower than that in the low-dose group (p= 0.0064) and PBS group (p= 0.0064).

There was a significant effect of drug treatment on the expression of Crh in the hippocampus (F(2, 21) = 8.699, P=0.0018) and amygdala (F(2, 21) = 3.643, P=0.0438). Post hoc analysis revealed that the mRNA expression of Crh in the high-dose group was significantly lower than that in the low-dose group in the hippocampus and amygdala (p= 0.0125, p=0.0135, respectively) and that the mRNA expression of Crh in the high-dose group was significantly lower than that in the PBS group (p= 0.0125) in the hippocampus.

There was a significant effect of drug treatment on the expression of Bdnf in the hippocampus (F(2, 21) = 11.01, P=0.0005) and amygdala (F(2, 21) = 6.707, P=0.0056). Post hoc analysis revealed that in the hippocampus, the mRNA expression level of Bdnf was significantly lower in the low-dose (p=0.0002) and high-dose (p=0.0024) groups than in the PBS group. In the amygdala, the mRNA expression level of Bdnf was significantly lower in the high-dose (p=0.0024) group than in the PBS group (0.0015).

### Lasting effects of HCQ on the mRNA expression of Il-1β, CRF, and Bdnf in the hippocampus and amygdala

As shown in figure 3, HCQ persistently altered the mRNA expression levels of Bdnf, IL-1β, and Crh in the hippocampus and amygdala. One-way ANOVA indicated that there was a significant effect of drug treatment on the expression of IL-1β in the hippocampus (F(2, 19) = 20.48, P<0.0001) but not in the amygdala (F(2, 19) = 0.7470, P=0.4872). Post hoc analysis revealed that the mRNA expression of IL-1 β in the high-dose group and low-dose group was significantly lower than that in the PBS group (p<0.0001 for both).

There was a significant effect of drug treatment on the expression of Crh in the hippocampus (F(2, 20) = 7.582, P=0.0035) but not in the amygdala (F(2, 20) = 0.5704, P=0.5743). Post hoc analysis revealed that the mRNA expression of Crh in the high-dose group and low-dose group was significantly lower than that in the PBS group (p= 0.0011 and p= 0.0143, respectively).

There was a borderline significant effect of drug treatment on the expression of Bdnf in the hippocampus (F(2, 20) = 3.255, P=0.0597) and a significant effect of drug treatment on Bdnf expression in the amygdala (F(2, 20) = 4.590, P=0.0229). Post hoc analysis revealed that in the hippocampus, the mRNA expression level of Bdnf was lower in the low-dose (p= 0.0231) and high-dose (p= 0.0742) groups than in the PBS group. In the amygdala, the mRNA expression level of Bdnf was significantly lower in the high-dose (p= 0.0068) group than in the PBS group.

## Discussion

The present study found that short-term repeated administration of HCQ led to a significant and persistent increase in anxiety-like behaviors in healthy mice and had an effect on depression-like behavior in the short term. HCQ also caused a sustained decrease in the mRNA expression of Il-1beta, Crh and Bdnf in the hippocampus; however, in the amygdala, HCQ temporarily affected the expression of Crh and induced a lasting significant decrease in the expression of Bdnf. Some of these behaviors and molecular changes, including the changes in anxiety-like behavior, the expression of Crh in the hippocampus and the expression of Bdnf in the amygdala, were dose-dependent, as the changes in the high-dose group were greater than those in the low-dose group. To the best of our knowledge, this study is the first to test the psychiatric effect of HCQ in an animal model, and our results suggest that, in healthy mice, HCQ administration can induce persistent behavioral changes and disrupt gene expression in the brain.

The HCQ treatment regimen used in this study mimics the current short-term regimen used for the treatment of SARS-COV-2. According to current clinical studies, most patients take HCQ for 5 to 7 days at a dose of 200 mg to 1200 mg on the first day followed by half of the original dose on the remaining days; in most studies, doses from 400 mg to 800 mg are administered on the first day. In this study, mice were given HCQ at one of two doses for 7 days. HCQ was administered at a dose of 164 mg/kg or 82 mg/kg on the first day and at a dose of 82 mg/kg or 41 mg/kg once daily for the next 6 days; these doses corresponded to 800 mg or 400 mg on the first day followed by 400 mg or 200 mg per day on the subsequent days in humans. The drug dose for mice was calculated based on the average body surface area ratio of mice to humans (mouse/human=12.3) (23). It should be noted that for this calculation, the dose per kilogram for humans was calculated based on the global average body weight for adults, which is 60 kg; however, the body weight of many people who take the drug may be higher. Therefore, the calculated dose administered to the animals may be higher than the corresponding dose administered to humans. Because the purpose of this study was to determine the potential risk of this drug, we thought it was important to choose a slightly higher dose and longer period of administration.

In this study, HCQ was administered by intraperitoneal (i.p.) injection because approximately 70%~80% of HCQ can be absorbed upon oral administration (13); thus, there would not be a large difference in drug absorption when the drug is administered orally and by i.p. injection. However, the speed of drug entry into tissues may be faster following i.p. injection than following oral administration. A previous study also indicated that the tolerated dose of i.p. injected HCQ is lower than that of orally administered HCQ (16); thus, we chose to administer the drug in two injections per day (one injection every 12 h). The body weight data showed that drug tolerance was basically good; except for a dose-dependent decrease in body weight on the first day, the rate of increase in body weight was not significantly different between the HCQ groups and the vehicle group. Additionally, evaluation of the locomotor activity of the animals did not show a significant difference between groups except for a HCQ-induced decrease in the locomotor activity of mice in the lasting group during the first 5 min of the test, which may have been a consequence of the anxiety-induced decrease in exploration in these animals.

Two time points were examined in this study, i.e., 24 h and 10 days after administration, and animal tissues were extracted 24 h after the completion of all the behavioral tests; thus tissues were collected 4 and 14 days after administration. Previous studies have shown that HCQ persists for a long time in vivo, with a half-life of 10-20 days in humans (13); however in rats, 80% of HCQ is eliminated by the 8th day, and 90% is cleared after 15 days (16). The metabolic rate of mice is approximately twice that of rats and 12 times that of humans (23), indicating that the direct effect of HCQ in the lasting group (10 days after administration) was likely quite small. Therefore, these two time points can effectively distinguish the immediate and lasting effects of HCQ.

In the current study, anxiety-like behavior was measured by using the EPM test, depression-like behavior was measured by using the FST, and working memory was measured by using the NOR test. The results showed that HCQ had a significant and sustained effect on anxiety, as indicated by decreased exploration time in the open arms. Depression-like behavior was also affected in the immediate group, as indicated by a significantly longer immobility time in the high-dose group than in the low-dose group but not in the vehicle group; these results suggest that HCQ has an effect on depression-like behavior. This increase in depression-like behavior is consistent with the changes in the expression of Crh in the amygdala. Our results did not reveal a significant effect of HCQ treatment on working memory, but total exploration time was decreased after HCQ administration, which also suggests that the drugs increased anxiety. Previous human studies have reported that HCQ can induce anxiety- and depression symptoms (10, 11), but no reports have shown that it impairs cognitive function. However, there have been only a few reports in humans, and there is a close relationship between cognitive impairment and psychiatric disorders (24). Furthermore, while we only measured working memory, the proteins in the brain that were affected by HCQ, such as Bdnf, have been indicated to be closely related to cognitive functions (25); therefore, our results cannot rule out the possibilty that HCQ affects cognitive functions.

The mRNA expression of three genes, namely, Il-1β, Crh and Bdnf, was measured in this study; these three genes are related to neuroinflammation, the HPA axis and neuroplasticity, respectively, and all of them are closely associated with the pathogenic mechanisms of mental illness (17, 26, 27). Our results showed that HCQ has a significant effect on all three molecules. At both time points, HCQ significantly decreased the expression of IL-1β in the hippocampus but had little effect on IL-1β in the amygdala, possibly because the hippocampus is more susceptible to changes in peripheral cytokine levels, as it is closer to the choroid plexus than the amygdala (28). Moreover, HCQ significantly decreased the expression of Crh in the hippocampus at both time points in a dose-dependent manner. Furthermore, HCQ had little effect on the expression of Crh in the amygdala; a significant effect was only observed in the immediate group, but the difference was mainly between the high-dose group and the low-dose group. In addition, HCQ significantly decreased the expression of Bdnf in both brain regions at both time points. It has been reported that these genes interact with one another; for example, it was reported that Il-1β can regulate the expression of Crh and Bdnf (29, 30). However, the widespread influence of HCQ on these genes is still beyond our initial expectation. Considering the importance and extensive influence of these genes on the CNS, the influence of HCQ on the CNS may be quite serious. Additionally, HCQ has a much longer half-life in humans than in rodents, and our results suggest that HCQ has significant effects on anxiety and gene expression more than 10 days after administration. Moreover, based on the expression pattern of IL-1β, the effect of HCQ on IL-1β increases after drug withdrawal. This implies that the influence of HCQ on human emotions and CNS functions may last for months. Previous case reports have also indicated that stopping drug administration cannot resolve its psychiatric effect quickly (31).

It should be noted that, as a preclinical study, the present study cannot prove that the clinical use of HCQ induces anxiety or depression in humans. The results of this study also do not clarify the mechanisms underlying the effect of HCQ on behavior or brain function. Based on these preliminary results, we speculate that the effect of HCQ on the CNS is mainly due to its peripheral immunosuppressive effects, which induce hypo-neuroinflammation in the brain, impairing the functions of immune cells in the brain, such as microglia. If our hypothesis is true, healthy individuals and patients with mild disease who do not suffer from infection-induced hyperinflammation would be at a higher risk of psychiatric symptoms than patients with severe SARS-COV-2. However, much more work is needed to prove this speculation. Our results prove that HCQ poses a serious risk to mental health. Given the current pandemic, we hope that based on the results reported in this study, scientists and clinical doctors will pay more attention to this potential adverse effect of HCQ and start to monitor the mental health status of patients in clinical trials. Furthermore, we suggest that individuals who are susceptible to psychiatric disorders should not take HCQ, especially as a prophylactic drug for SARS-COV-2.

